# Dietary lipids modulate Notch signaling and influence adult intestinal development and metabolism in *Drosophila*

**DOI:** 10.1101/273813

**Authors:** Rebecca Obniski, Matthew Sieber, Allan C. Spradling

## Abstract

Tissue homeostasis is a complex balance of developmental signals and environmental cues that dictate stem cell function. However, it remains poorly understood how nutrients interface with developmental pathways. Using the Drosophila midgut as a model we found that during the first four days of adult life, dietary lipids including cholesterol, determine how many enteroendocrine (ee) cells differentiate and persist in the posterior midgut where lipids are preferentially absorbed. The nuclear hormone receptor Hr96 which functions to control sterol trafficking, storage, and utilization, is required for sterol-mediated changes in ee number. Dietary cholesterol influences new intestinal epithelial cell differentiation from stem cells by altering the level and persistance of Notch signaling. Exogenous lipids modulate signaling by changing the stability of the Delta ligand and Notch intracellular domain and their trafficking in endosomal vesicles. Lipid-modulated Notch signaling occurs in other nutrient-dependent tissues such as the ovary, suggesting that Delta trafficking in many cells is sensitive to cellular sterol levels. These diet-mediated alterations in ee number in young animals contribute to a metabolic program adapted to the prevailing nutrient environment that persists after the diet changes. A low sterol diet also slows the proliferation of enteroendocrine tumors initiated by disruptions in the Notch pathway. These studies show that a specific dietary nutrient can modify a key intercellular signaling pathway to shift stem cell differentiation and cause lasting changes in tissue structure and physiology.

## Introduction

Development is a unique partnership between a flexible genetic program and the environment. Major advances have occurred in recent years in understanding the genes and pathways that program embryo and tissue development, many of which are highly conserved among diverse invertebrate and vertebrate animals. In addition, numerous examples have been documented where environmental variables, nutrition in particular, exert or are believed to exert strong modulating effects on developmental outcomes and disease susceptibility (reviewed in Preston et al. 2018). Examples, include folate and neural tube closure (Imbard et al. 2013), cholesterol and heart disease (Orho-Melander, 2015), and caloric intake and longevity (Templeman and Murphy, 2017). Despite the great interest and importance of understanding how the environment modifies development, gaining a mechanistic understanding of these unprogrammed effects on animal form and function has proved much more difficult than deciphering the developmental program itself.

Diet is one of the most important and ubiquitous environmental variables impacting animal life cycles. A nutrient poor diet during embryonic development may establish a scarcity metabolic program in the offspring that can lead to metabolic syndrome (review: Ramakrishnan, 2004). Dietary intake of certain nutrients may also increase the risk of cancer. For example, there is a strong correlation between a high fat diet and colon cancer (Beyaz et al., 2016; O’Neill et al., 2016) raising the possibility that a high fat diet may influence the developmental signaling pathways that promote cell proliferation. However, nutrients such as cholesterol affect cancer incidence in many potential ways, complicating our understanding of the connections between dietary cholesterol intake and oncogenesis (Kloudova et al. 2017).

Cholesterol uptake and utilization is controlled by mechanisms that are highly conserved between Drosophila and mammals. For example, intestinal enterocytes in both groups take up dietary cholesterol by endocytosis and vesicle trafficking into lysosomes. Subsequently, under control of Niemann-Pick type C genes NPC1 and NPC2, cholesterol is moved from lysosomes to other organelles such as endoplasmic reticulum (ER) and mitochondria (Huang et al 2005). Once in the ER a large portion of the cholesterol is esterified by acetylCoA cholesterol acyltransferase (ACAT), packaged into lipoprotein particles and transported to peripheral tissues. In mammals, cholesterol biosynthesis is controlled by the lipid-mediated regulation of protein stability in the endoplasmic reticulum (Goldstein and Brown, 1990). When cholesterol levels in enterocytes are high, the rate limiting biosynthetic enzyme, the membrane protein HMGCoA reductase, is destabilized by ubiquitination, dislocated out of the ER and degraded via the endoplasmic reticulum-associated degradation (ERAD) pathway. In contrast, in lipid-deprived cells, Insig-1, a key protein involved in HMGCoA turnover and in processing of sterol regulatory element binding proteins (SREBPs), is degraded via ERAD. Even though *Drosophila* is a cholesterol auxotroph, mammalian lipid regulatory proteins are still subject to sterol-regulated ERAD showing that lipid mediated control of protein stability in the ER via ERAD has been highly conserved (Carvalho et al. 2006; Faulkner et al. 2013).

The expression of genes mediating cholesterol uptake, as with many other dietary lipids, is mediated largely through nuclear receptors (NR). These ligand-regulated transcription factors recognize small lipophilic molecules and mediate broad changes in gene expression to regulate development, homeostasis, and metabolism (Evans and Mangelsdorf, 2014). Cholesterol efflux in mammals is regulated primarily by NR subfamily 1 members LXRα and LXRβ. In response to high intracellular oxysterols, LXR induces the expression of cholesterol transport proteins thereby stimulating cholesterol trafficking from the peripheral tissues to the liver where it is eliminated in the form of bile (Kalaany and Mangelsdorf, 2006). In *Drosophila,* a single subfamily 1 ortholog, Hr96, is expressed in the intestine and the fat body, and is required for development in cholesterol-depleted environments (Horner et al., 2009). Consequently, NR1 members and their conserved target genes are strong candidates for mediating cholesterol-induced effects on intestinal development.

The *Drosophila* midgut is powerful model system in which to analyze the relationship between diet and intestinal development at the cellular, molecular and functional levels, in part because nutrient processing is highly regionalized (Buchon et al., 2013; Marianes and Spradling, 2013). The activity within each gut subregion is strongly impacted by distinct self-renewing intestinal stem cells (ISCs) (Micchelli and Perrimon, 2006; Ohlstein and Spradling, 2006) whose daughter cells differentiate in response to a high Notch signal into nutrient-processing enterocytes (ECs), or in the presence of low intracellular Notch activity into multiple types of hormone-secreting enteroendocrine cells (ee’s) (Ohlstein and Sprading, 2007; Guo and Ohlstein, 2015). Notch signaling is largely regulated by its ligand, the transmembrane protein Delta, whose activity is influenced by factors that impact the trafficking, recycling and turnover within endosomal vesicles of both Delta and Notch itself (Seugnet et al., 1997; Conner, 2016). In the mammalian intestine, Notch signaling also specifies enterocytes and enteroendocrine cells (Demitrack and Samuelson, 2016). Abnormalities in Notch signaling are associated with increased cell proliferation in both *Drosophila* and mammals, including colon cancer (Suman et al., 2014).

We have found that dietary cholesterol influences intercellular Notch signaling in the adult midgut. When cholesterol is high, Delta levels and Notch signaling are relatively low which favors the differentiation of enteroendocrine cells. When cholesterol is low, Delta accumulates to high levels in intercellular vesicles, Notch signaling is stimulated and enteroendocrine cell differentiation is reduced. These effects of dietary sterols require the nuclear hormone receptor, Hr96, which controls genes, including Npc2b, that mediate sterol trafficking, storage, and utilization. The consequences of dietary lipids are particularly dramatic during the first four days of adulthood, when many enteroendocrine cells normally develop in posterior regions that are specialized for lipid absorption. The number of posterior enteroendocrine cells that arise during this time is strongly influenced by lipid levels in the early diet. These cellular differences persist and appear to provide a metabolic memory, as flies raised initially under lipid deprivation later in life accumulate 18% more sterols on a standard diet than controls. Moreover, dietary lipid levels affect the progression of enteroendocrine tumors. Hr96 similarly influenced Delta levels and Notch signaling in another nutrient-dependent tissue, the ovary, and in mammalian cells. Thus, our results reveal a specific and potentially widespread mechanism through which a dietary nutrient affects Notch intercellular signaling and cellular differentiation with consequences for adult metabolic programming and cancer susceptibility.

## Results

### Dietary sterols influence adult midgut enteroendocrine cell number

The *Drosophila* midgut completes development shortly after adult eclosion, like the mammalian intestine which develops rapidly after birth. Delayed differentiation is proposed to allow intestinal metabolism to adapt to an individual’s nutrient environment (reviewed in Reynolds et al. 2015). We observed a delayed appearance of intestinal SREBP signaling (Figure 1A,B), a key pathway in the regulation of lipogenesis (Brown and Goldstein, 1997; Kunte et al., 2006). Only sparse, low level activity is present in the intestine of newly eclosed flies (Figure 1A), consistent with previous studies (Reiff et al., 2015). In contrast, the pathway is highly activated in the posterior midgut of 5-day old flies (Figure 1B). Consequently, we investigated whether midgut differentiation in young adults is influenced by the lipid content of their initial diet. We collected late stage pupae and allowed these animals to eclose and feed on either a control or lipid-depleted diet for 10 days. We then dissected these flies and stained both the anterior and posterior midgut with markers to assess their cellular composition (Figure 1C).

**Figure 1:**
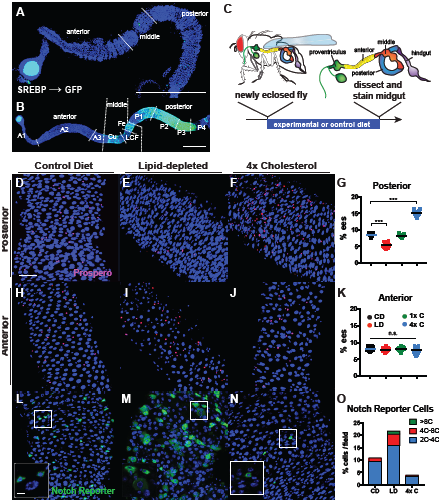
Dietary sterols influence adult midgut enteroendocrine cell number. (A-B) Midguts from 6-12 hour old (A) or 5 day old (B) adult females. SREBP reporter (green) DAPI (blue). Scale bar = 400µm (C) Experimental strategy to test the effects of diet on differentiation. (D-K) Posterior (D-G) or Anterior (H-K) midgut regions from animals fed a control diet (CD) (D,H), lipid-depleted diet (LD) (E, I), or lipid-depleted diet supplemented with 4x cholesterol (4X C) (F, J). DAPI (blue); Prospero (red). Scale bar = 40µm. (G, K) % enteroendocrine (ee) cells (Prospero^+^) relative to total posterior (G) or anterior (K) cells. Circles show individual values color-coded for diet as indicated (N = 20 for all samples). Bar = Average value; ^***^ = p < 0.005 (t-test). (L-O) Notch signaling reporter Su(H)GBE-GAL4 driving UAS-CD8-GFP in posterior midgut. Animals were fed on control(L), lipid depleted (M) or 4X cholesterol-supplemented diet (N), for 10 days prior to analysis. DAPI (blue); GFP (green). Inset scale bar = 10µm. (O) % of total cells with Notch signaling (GFP^+^) in a 500µm^2^field. GFP^+^ cells were subdivided by ploidy based on nuclear size and DAPI intensity: blue (< 4C), red (4C-8C), green (>8C).

When we looked at the distribution of midgut cell types in adult animals fed a lipid-depleted diet, we observed significant decrease in posterior enteroendocrine (ee) cells (recognized by Prospero staining) relative to controls (Figure 1D,E). This reduction was not observed in the anterior midgut (Figure 1H,I). To confirm that the absence of dietary lipids caused the reduction in ee number, we fed newly eclosed flies a lipid-depleted diet supplemented with a single lipid species in the form of cholesterol, fatty acid, or triglyceride, and found that providing these dietary metabolites prevented the reduction in ee number (Figure1G, Figure S1E-H). The effect was specific to the period of initial adult gut development, because flies shifted to a lipid depleted diet for 10 days after one week on control media did not show a significant change in ee number (Figure S1D).

To determine whether elevated dietary lipid levels could likewise affect ee number, we supplemented a lipid-depleted diet with high levels of specific lipid species. Increasing the concentration of dietary fatty acid resulted in death within a week of feeding, however, animals fed a high cholesterol diet are viable and fertile. In newly eclosed animals fed a diet with four times the amount of cholesterol in a control lab diet the frequency of posterior ee cells was significantly increased (Figure 1F). When cholesterol supplementation commenced one week after eclosion, this increase did not occur, even after 14 days of high cholesterol (Figure S1D). We found that, as expected, growth on a lipid-depleted or 4X cholesterol diet significantly changed the levels of intestinal cholesterol and cholesterol ester (Figure S1C). Since *Drosophila* are sterol auxotrophs that normally acquire sterols from yeast rather than cholesterol, we fed newly eclosed flies on lipid-depleted media supplemented with 4x ergosterol, the major yeast sterol and observed a similar increase in the percentage of enteroendocrine cells in the posterior midgut (Figure S1G,H). Cholesterol supplementation or depletion does not act by altering rates of ISC division, because the effects of these treatments on the mitotic index (ISC divisions) did not correlate with ee production (Figure S1M). Furthermore, ee’s did not change in frequency due to altered death rates, because the number of TUNEL-positive cells per midgut was unchanged following cholesterol supplementation and cell death increased rather than decreased following lipid depletion (Figure S1N).

Notch signaling regulates the differentiation of ISCs into enterocytes or enteroendocrine cells. To determine whether Notch pathway activation is affected by dietary lipids, we used a reporter for Notch signaling, GBE-Su(H)-GAL4, and found that animals fed a low cholesterol diet had significantly more midgut Notch signaling activity than animals fed a control diet (Figure 1L,M). Moreover, a lipid depleted diet caused Notch signaling activity in enteroblasts to remain high as they developed into enterocytes as evidenced by poyploidization (Figure 1M,O). In contrast, animals fed a 4X cholesterol-supplemented diet had significantly fewer cells with Notch reporter activity (Figure 1N,O). This suggests that increasing or reducing dietary cholesterol levels decreases or increases the level and kinetics of Notch signaling, and hence increases or reduces enteroendocrine cell production, respectively.

### Hr96 mediates cholesterol-dependent control of enteroendocrine cell number

Physiological effects of dietary cholesterol on enteroendocrine cell number would require cholesterol uptake and cellular utilization, pathways that depend on the nuclear hormone receptor Hr96 (Horner et al., 2009). Consequently, we analyzed *Hr96*^*1*^ animals, which lack all Hr96 activity (King-Jones et al., 2006), fed a control diet. *Hr96*^*1*^ midguts contained significantly fewer ee’s compared to heterozygous controls in the posterior region (Figure 2A,B), and throughout the midgut (Figure S2A-H). These reductions strongly resemble the effects of a lipid-depleted diet on wild type animals, suggesting that without normal lipid uptake and processing under the control of *Hr96*, animals are effectively lipid-starved.

**Figure 2:**
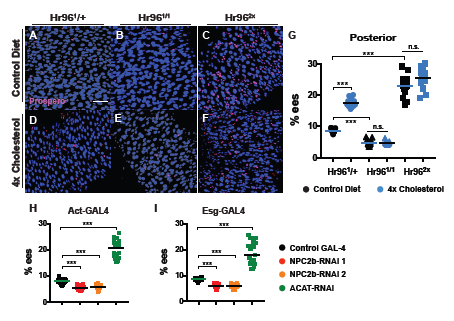
*Hr96* mediates sterol availability and enteroendocrine cell number. A-F) Poterior midguts from animals of the indicated genotypes following culture on the indicated diets. (A,D) *Hr96*^*1*^/ + (as a control); B,E) *Hr96*^*1/1*^; C,F) *Hr96*^*2X*^ (duplication). Animals were fed a control diet (A-C) or 4X cholesterol-supplemented diet (D-F) for 10 days prior to analysis. DAPI (blue); Prospero (red); scale bar = 40µm (G) % ee cells relative to total posterior cells. Data points show individual values shape-coded for genotype and color-coded for diet as indicated (N = 15 for all samples). Bar = Average value; ^***^ = p < 0.005 (t-test), n.s. = not significant. (H, I) The *Hr96* target gene *NPC2b*, and *ACAT* have opposite effects on ee differentiation. % ee cells relative to total midgut posterior cells for animals expressing the indicated RNAi constructs driven in enterocytes by Act-GAL4 (H) or in ISCs and enteroblasts by esg-GAL4 (I) and raised on a control diet.

We investigated whether the normal gene dosage of *Hr96* is rate-limiting for sterol utilization by examining midguts from animals carrying two tandem copies of *Hr96* on each homolog (*Hr96*^*2X*^) (Sieber and Thummel, 2009). *Hr96*^*2X*^ flies contained significantly more enteroendocrine cells in all regions of the intestine (Figure 2C; Figure S2C,G). Interestingly, on a high sterol diet *Hr96*^*2X*^ animals did not significantly increase ee’s beyond the high level present on a control diet (Figure 2F,G). Thus, increased *Hr96* dosage is sufficient on a control diet to utilize enough sterol to achieve a maximal effect on ee number. Expressing *Hr96* only in enterocytes using *Myo1A-GAL4* was sufficient to increase ee number relative to controls (Figure S2I-K). This suggests that dietary sterols can be taken up and processed by differentiated enterocytes before they are transferred to ISCs where they can influence Notch signaling and cell differentiation.

*Drosophila* has two NP1 class C genes, and eight NP2 class C genes, and several of these genes are regulated by Hr96 (Horner et al. 2009). These include *Npc1b* which is required for dietary cholesterol absorption (Voght et al. 2005), and at least two *Npc2* class genes involved in intracellular cholesterol trafficking, *Npc2a* and *Npc2b* (Huang et al., 2005; 2007). To examine whether *Hr96* acts on ee number by controlling sterol transport genes, we used two different RNAi lines to suppress the *Hr96* target gene *Npc2b* in the adult flies. The posterior midguts of animals expressing either RNAi construct contained fewer ee’s compared to controls (Figure 2H; Figure S2L-M). This was similar to the behavior of animals on a lipid-depleted diet, consistent with the predicted inability of *Npc2b* mutant animals to take up and utilize sterols. Interestingly, knocking down *Npc2b* only in stem cells and undifferentiated cells using esg-GAL4 in the intestine still reduced ee number (Figure 2I; Figure S2L-R). Thus, cholesterol transport and availability in undifferentiated cells as well as in enterocytes influences the differentiation of intestinal stem cell daughters.

Another important and conserved gene required for steroid metabolism in mammals and *Drosophila* is ACAT, which mediates sterol esterification, transport and storage. We used RNAi to inhibit the Drosophila ACAT homolog, CG8112 (Horner et al. 2009), and found a subsequent increase in the percentage of ee’s in the midgut, similar to the effects of high dietary cholesterol (Figure 2H,I, Figure S2N,Q). This is consistent with the expectation that ACAT knock-down will increase free intracellular sterols (Temel et al., 2007). These experiments strongly support the view that the *Hr96* and *Npc2b*-dependent uptake of dietary sterols in the midgut influences the differentiation of enteroendocrine cells in young adults.

### RNAseq analysis of diet-dependent midgut gene expression

The *Drosophila* midgut expresses distinctive gene programs along its anterior-posterior axis within multiple tissue subregions (Marianes et al. 2013; Buchon et all. 2013). To investigate the effects of dietary lipids on midgut gene expression, we performed RNAseq in triplicate on anterior and posterior midguts from wild type animals fed a control, lipid-depleted, or cholesterol-supplemented diet. Parallel studies were done on anterior and posterior midguts from *Hr96*^*1*^ and *Hr96*^*2X*^ mutants. All 30 samples were aligned to the release 6 genome, and analyzed using a standard pipeline involving TopHat, Cufflinks, and CuffDiff as described in Methods. General characteristics of the significant expression differences between these conditions are summarized in Figure S3A-C.

These data provide insight into the nature of the additional enteroendocrine cells that are induced by supplemental dietary cholesterol and by *Hr96* duplication. Since *Hr96*^*2X*^ animals contain the largest increase in enteroendocrine cells, approximately three-fold in the posterior, we examined the expression levels of known enteroendocrine hormone genes in *Hr96*^*2X*^ midguts to query which ee subtypes were likely to have increased. Six of the seven enteroendocrine hormone genes known to be expressed in posterior ee cells (Marianes et al. 2013; Veenstra and Ida, 2014) were significantly increased in posterior midguts from *Hr96*^*2X*^ animals compared to controls (Figure 3A). The bHLH transcription factor Dimmed, which mediates peptide secretion via the large dense core vesicle pathway used by ee cells (Hamanaka et al. 2010; Beehler-Evans et al. 2015), was also significantly increased (1.27 +/-0.18 vs 0.76 +/-0.26 p<0.05, t-test) in *Hr96*^*2X*^ posterior midguts, consistent with an increased number of functional, neuropeptide-secreting ee cells. These results argue that the increase in enteroendocrine cells in response to dietary sterols involves most or all of the ee cell types in the posterior region, rather than a specific response to a dietary molecule by a single ee cell type. Two enteroendocrine hormone genes, AstC and Orcokinin, increased significantly in expression in the anterior midgut of *Hr96*^*2X*^ animals (Figure 3B). Changes in anterior hormone expression are smaller than those in the posterior, consistent with the smaller observed increase in anterior ee cell number.

**Figure 3:**
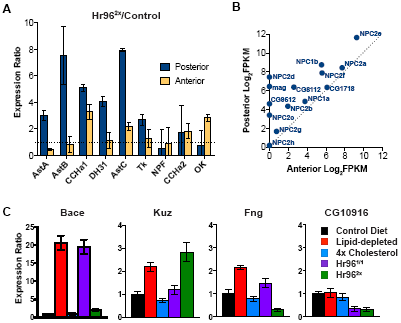
Dietary lipids differentially alter gene expression in anterior and posterior midgut. (A) The relative expression in anterior or posterior midguts of the nine known enteroendocrine peptide hormone genes from *Hr96*^*2X*^ animals (elevated ee number) compared to controls. (B) Expression (log2fpkm) of the indicated genes implicated in lipid metabolism in posterior (y-axis) vs anterior (x-axis) midgut. (C) Expression ratio in posterior midgut relative to control for indicated genes implicated in Notch signaling under indicated dietary conditions, or in different genotypes on a control diet

The RNAseq data provide further evidence that sterol absorption takes place preferentially in the posterior midgut (Figure 3B; Marianes et al. 2013). Genes involved in sterol uptake and utilization, including the ten NPC class genes, ACAT and 3 other cholesterol-induced genes: magro, CG32186, CG11425 (wunen-like), Lip3, Nlaz, were expressed at higher levels in the posterior compared to the anterior midguts of animals fed a control diet. Only a few of these genes were strongly reduced in Hr96 mutants (Npc2d, Npc2f, mag, CG42816, Lip3). This is not surprising, however, as NPC genes involved in Hr96-regulated cholesterol uptake have other functions, for example in gut immunity (Shi et al. 2012). Hr96 RNA was significantly increased in response to lipid depletion: posterior midgut: 157+/-19.6 vs 96.4+/-37.8; p<0.05 t-test; anterior midgut: 152.8+/-12.8 vs 55.8+/-12.0 p<0.001 t-test.

### Dietary sterols modulate Delta levels and trafficking

Dietary lipids, and cholesterol in particular, influence midgut cell differentiation and modulate Notch signaling and at a time when the posterior subregions are completing their development. To investigate how dietary lipids alter Notch signaling, we immunostained the Notch ligand Delta to visualize its expression in the posterior midguts of 10-day old animals fed various diets. Delta was found in cytoplasmic vesicles in control animals within scattered diploid cells that correspond to ISCs (Figure S4A) as previously reported (Ohlstein and Spradling, 2006). ISCs from animals fed a lipid-depleted diet consistently stained more strongly for Delta compared to controls (Figure S4B), Moreover, in these animals Delta staining could be readily detected within daughter enteroblasts generating pairs of Delta-labeled cells (Figure S4B, B’). Since posterior midgut size did not increase as a result of lipid depletion, these cell pairs were likely caused by reduced Delta turnover in enteroblasts under lipid-deprived conditions rather than stem cell amplification divisions (O’Brien et al. 2011; Buchon et al. 2010). Conversely, feeding young animals a high sterol diet, reduced the level of Delta staining within ISCs (Figure S4C).

We examined Delta levels in animals mutant for cholesterol uptake genes to verify that these changes in Delta content depended on physiological lipid uptake. *Hr96*^*1*^ mutants on a normal diet showed elevated levels of Delta in ISCs compared to controls (Figure 4A,B). When these animals were grown on lipid depleted media, the levels of Delta were further increased and abundant Delta-rich cytoplasmic vesicles were now readily apparent (Figure 4 E,F). In contrast, Delta levels were scarcely detectable and foci were rare in *Hr96*^*2X*^ animals, even when fed a lipid-depleted diet (Figure 4C,G). *NPC2b* mutant animals, which like *Hr96*^*1*^ mutants are defective in cholesterol incorporation and mimic wild type animals on a lipid-depleted diet, showed higher levels of cytoplasmic Delta staining compared to control (Figure S4E-G). In both *Hr96*^*1*^ and *Npc2b* RNAi animals, Delta protein expression persisted in enteroblasts and young enterocytes (Figure 4F, Figure S4F,G), like control animals on a lipid-depleted diet (Figure 4B’).

**Figure 4:**
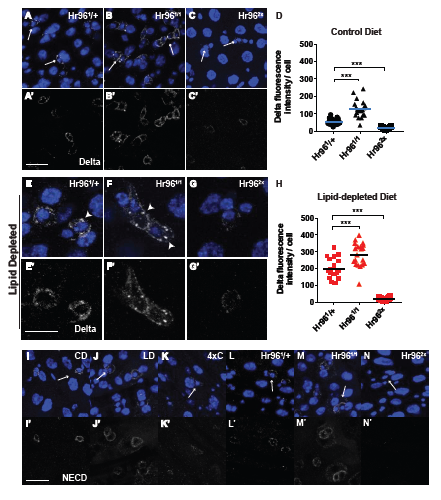
Hr96-mediated sterol metabolism regulates Delta protein levels and trafficking. Posterior midguts are shown from (A,E) *Hr96*^1/+^ (B,F) *Hr96*^1/1^and (C,G) *Hr96*^*2X*^ animals fed a control diet (A-C) or a lipid depleted diet (E-G) for 10 days prior to analysis. DAPI (blue) and Delta (white). arrows: ISCs (A, C), ISC-enteroblast pairs (B). (D,H) Corrected fluorescence intensity per Delta^+^ cell is plotted in arbitrary units from animals of the indicted genotypes. Individual values are shape-coded for genotype as indicated. N = 15 (D); N = 20 (H). Bar = Average value; ^***^ = p<0.005; (t-test). Scale bar = 5µm for all images; I-N) Notch extracellular domain (NECD) expression (white) is shown in posterior midguts of wild type animals raised on a control (I), lipid-depleted (J) or 4X cholesterol (K) diet, or from *Hr96*^1/+^ (L) *Hr96*^1/1^(M) or *Hr96*^*2X*^ (N) animals raised on a control diet.

Taken together, these observations show that when lipids including cholesterol are low due to diet or a genetic defect, ISCs continue to proliferate but turn over Delta more slowly than normal. This change in Delta persistence matches the increased Notch signaling activity previously observed on lipid-depleted diets (Figure 1M) and is likely to favor EC over ee production from ISCs. Conversely, *Hr96*^*2X*^ animals on a control diet (Figure 4C) or ACAT RNAi animals on a control diet (Figure S4H) showed reduced Delta expression. This is similar to wild type animals fed a high sterol diet (Figure S4C) and is consistent with their reduced Notch signaling activity (Figure 1N), a condition likely to favor ee over EC production. These data strongly suggest that gut lipid uptake and utilization influences endomembrane lipid composition, and that these changes influence Delta abundance through their effects on Delta subcellular localization, trafficking and turnover.

### Notch is also modulated by lipid uptake and utilization

We investigated whether the abundance and trafficking of Notch is also affected by dietary lipid utilization. Proper recycling of the Notch extracellular domain (NECD) is crucial for accurate Notch activation or suppression and is thought to couple to Delta turnover (Coumailleau et al., 2009). If lipid depletion impairs Delta endosomal recycling or degradation, we reasoned that the NECD may also display aberrant turnover and trafficking. Using antibodies against the NECD we examined the intestines of flies fed a lipid-depleted diet and of *Hr96* mutants, and found that NECD staining was present in putative ISC-EB cell pairs (Figure 4J,M). In contrast, NECD was only detectable in single cells in control animals (Figure 4I,L). Furthermore, NECD positive cells had less antibody staining in high sterol or Hr96 overexpression animals (Figure 4K,N). Taken together, these data further support the finding that cellular sterol levels strongly influence the trafficking and turnover of Notch pathway proteins, including Delta and Notch NECD.

The RNAseq experiments suggested several genes among those known to influence Notch signaling that might be responsible for these changes in protein stability (Figure 3C, Figure S3D). The membrane protease, Bace, a predicted Notch ligand regulator, is one of the most highly up-regulated genes in lipid depleted or *Hr96* mutant midguts. *Kuzbanian (kuz)*, an ADAM metalloprotease was significantly up-regulated (+2.3 fold) after lipid depletion and slightly decreased on a high sterol diet. *Kuz* mutant ISC clones have a *Notch* mutant phenotype, favoring ee production (Lucchetta and Ohlstein, 2017). The glycosyl transferase, Fringe (Fng) enhances Notch-Delta interaction and *fng* mRNA is significantly increased after lipid depletion. *CG10916* encodes a predicted E3 ligase that interacts with Notch (Guruharsha et al., 2011) and behaves as an *Hr96*-regulated gene. Degradation of the Notch intracellular domain (NICD) by the proteasome is important for turning off Notch transcriptional activity.

### Sterol regulation of Delta is also observed in the ovary and in human cells

To investigate whether the relationship between cellular sterol levels and Notch signaling described above for the Drosophila midgut also applies to other tissues, we examined the Drosophila ovary. Oogenesis is highly dependent on efficient nutrient processing, since proteins, lipids, and carbohydrates, all initially processed by the midgut, are accumulated in a tightly regulated, stepwise manner to support the growing oocyte (Sieber and Spradling, 2015). Like midgut enterocytes and enteroendocrine cells, the follicle cells of the ovary also rely on Notch signaling to induce follicle cell differentiation at the mitotic-endocycle (ME) transition, and to coordinate somatic and germline development (Sun and Deng, 2005). Diet strongly influences the rate of follicle development and passage through checkpoints (Drummond-Barbosa and Spradling, 2001), so we used genetic manipulation of sterol metabolism to look for connections to Notch signaling.

Delta was present on membranes at only low levels in wild type mid-stage follicles (Figure 5A), consistent with previous reports (Bender et al., 1993). However, significantly more Delta was present on membranes from similarly staged *Hr96*^*1*^ mutant follicles (Figure 5B). Furthermore, large cytoplasmic Delta foci were observed in *Hr96*^*1*^ mutant follicles (Figure 5D), but not in controls (Figure 5C), reminiscent of the Delta foci in midgut ISCs and developing enterocytes. Similarly, we found *Hr96*^*1*^ ovaries displayed increased levels of NECD (Figure 5F) relative to controls (Figure 5E). Large NECD aggregates were present in stage 9-10 follicles of *Hr96*^*1*^ mutants (Figure 5F), but not controls (Figure 5E). Corresponding increases in membrane Delta and NECD were observed in ovaries from NPC2b-RNAi animals as well (Figure S5A-E). To determine whether these effects on protein levels and aggregation were specific to components of the Notch pathway, or affected other transmembrane proteins, we stained control and *Hr96*^*1*^ mutant follicles with an antibody against the LDLR homolog LpR2 and found no significant change in LpR2 levels or localization (Figure 5G, H). These observations argue that sterol levels have similar effects on Notch signaling components in ovarian cells as in the midgut.

**Figure 5:**
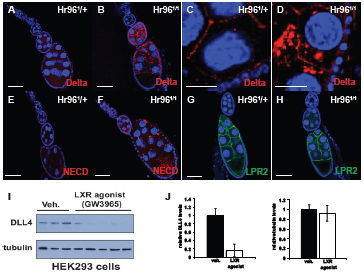
Delta and NECD levels and trafficking are specifically modulated by lipids in the ovary and in mammalian cells. Notch (A-D), NECD (E-F), or Lpr2 (G,H) immuofluorescence is shown in *Drosophila* ovarian follicles from *Hr96*^1/+^ (A,C,E,G) or *Hr96*^1/1^(B,D,F,H). C and D show expression in the cytoplasm of a large polyploid nurse cell. DAPI (blue) Scale bar = 50?m. (I) Western blot assaying Delta-like ligand 4 (DLL4) levels from HEK293 cells treated with vehicle alone (Veh.) or LXR agonist. Tubulin staining (below) served as a loading control. Lanes are from independent assays. J) Average DLL4 levels and tubulin levels (bars = +/-SD) were plotted as determined from the blot in (I).

Notch signaling regulates the ME transition in follicle cells during stage 5-6 of oogenesis (Sun and Deng, 2005; 2007). Late stage follicles in *Hr96*^*1*^ mutants had significantly more follicle cells than similarly staged controls (Figure S5F,G,J), indicating that the Notch-induced ME transition had been delayed. Moreover, *Hr96*^*1*^-mutant stage 14 follicles contained follicle cell nuclei that had been extruded from the follicular layer, further suggesting that a delayed or incomplete switch to the endocycle resulted in excess follicle cells (Figure S5H,I,K). Normally, the Notch-regulated gene *hindsight* (*hnt)* is expressed in follicle cells after the ME transition (Figure S5L). In contrast, Hr96 mutants expressed Hnt slightly earlier than normal, but Hnt expression levels remained lower than in control follicles (Figure S5M) suggesting that altered Notch signaling ultimately delayed rather than accelerated normal follicle development. This defect in Notch signaling may stem from the fact other sterol dependent pathways such as ecdysone signaling also function to regulate the M/E transition and defective cholesterol metabolism may cause disruptions in multiple pathways in the ovary.

Since all the components linking intracellular sterol levels to Notch signaling are highly conserved between *Drosophila* and mammals, we investigated whether human cells show similar responses. We identified a human cell line that expresses the cholesterol regulator LXR, as well as Delta-like Notch ligand 4 (DLL4) and asked whether promoting intracellular reverse cholesterol transport with the LXR agonist GW3965 would affect DLL4 protein levels. DLL4 protein levels were consistent in cells treated with vehicle alone, but after treatment with the LXR agonist, DLL4 protein decreased significantly in 5 independent experiments (Figure 5I,J). This result suggests that intracellular sterol levels act in a conserved manner on Notch ligand stability and Delta-Notch signaling in diverse tissues and organisms.

### Early lipid depletion reduces posterior ee numbers and establishes a scarcity metabolic state

Our observation that dietary lipids and cholesterol in particular enhances Notch signaling in the midgut and ovary raises the question of how such a connection affects the animal phenotype. For example, dietary lipids modulate the final differentiation of the gut in young animals but do changes in ee number and potentially other characteristics persist and influence the animal’s response to a different dietary regime? To determine whether nutrients in the early diet influences metabolism after the diet has changed, we raised young flies on lipid-depleted food feed for one week to increase Notch signaling and decrease posterior ee number. We then transferred them the flies to normal yeast food and observed that the reduction in posterior ee number persisted for at least two weeks (Figure 6A). Interestingly, flies raised initially on lipid-depleted media, after one week back on normal yeast food, accumulated 18% more whole body cholesterol per fly than animals that had eaten a cholesterol-rich diet beginning at eclosion (Figure 6B). This suggests that low lipid levels early in development generate a scarcity metabolic phenotype that parallels the reductions in ee number, and that programs greater lipid storage under normal dietary conditions. When tested two weeks after switching from a lipid-depleted to a normal diet, the animals still had fewer posterior ee cells and stored 14% more whole body cholesterol than controls that were never lipid-deprived (Figure 6A,B). Thus, the effects of nutrient starvation early in adult life can persist for a significant period in the absence of the original stimulus.

**Figure 6:**
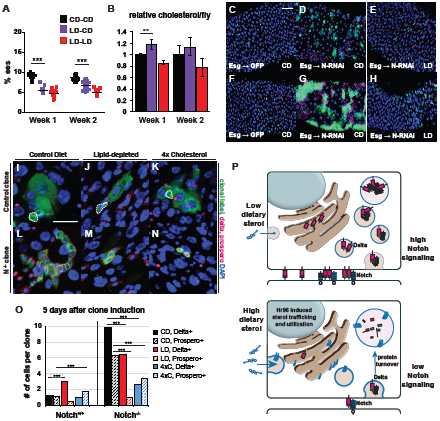
A low lipid diet generates a persistent scarcity metabolic state and delays enteroendocrine tumor formation. % ee’s in posterior midgut (A) is lower and relative total body cholesterol per fly (B) is greater in flies raised on a lipid-depleted diet for 7d and then shifted to control diet for 1 or 2 additional weeks (LD-CD: purple), than in flies raised continuously on a control diet (CD-CD, black). Flies raised and maintained on an LD diet (LD-LD:red) were measured for comparision. (N = 10) ^**^ = p<0.01, ^***^ = p<0.005 (t-test). (C-H) A low lipid diet suppresses enteroendocrine tumor production. Posterior midguts from control (Esg::GFP) or Notch-RNAi (Esg::N-RNAi) fed a control (CD) or lipid-depleted (LD) diet for 4 days (C-E)) or 9 days (F-H) after tumor initiation. DAPI (blue); GFP (green); Prospero (red, nuclear) and Delta (red, cytoplasmic). Scale bar = 40µm. (I-N) Posterior midguts from animals fed the indicated diets and showing MARCM clones 9 days after induction of control (I-K) or *N*^*-/-*^ cells (L-N) marked with GFP. Scale bar = 10µm. (O) Mean clone size from experiment in (I-N). ***p<0.005 (t-test). (P) Model of dietary sterol-mediated regulation of Notch signaling via changes in Delta and Notch trafficking and stability in the ER.

### Dietary lipids influence enteroendocrine tumor development

When Notch signaling is strongly reduced in ISCs they give rise to rapidly proliferating neoplasias composed of ISC-like and enteroendocrine-like progenitors (Micchelli and Perrimon, 2006; Ohlstein and Spradling, 2006; Patel et al., 2015). Strikingly, these “enteroendocrine tumors” preferentially form in lipid-rich regions of the intestine, which are mostly located in the posterior midgut (Marianes and Spradling, 2013). Our finding that elevated dietary sterol downregulates Notch signaling suggests an explanation for the preferential development of enteroendocrine tumors in lipid-rich midgut regions. ISCs in lipid-rich zones would likely contain elevated levels of membrane cholesterol and other lipids that would further depress Notch signaling by the mechanism described here, enhancing the effect of genetic lesions in the Notch pathway. Thus, tumor formation might represent another adult characteristic affected by diet.

If increased dietary lipids suppress Notch signaling in stem cells and thereby promote enteroendocrine tumors, we reasoned that a lipid-depleted diet, which increases Notch signaling, might slow tumor development. We induced tumor formation by expressing Notch RNAi in progenitor cells throughout the intestine using esg-GAL4, and tracked tumor development quantitatively by counting the number of clustered tumor cells that were labeled using UAS-GFP and Prospero staining. Four days after expressing N-RNAi, at least 80% of midguts from animals on control diets showed multiple tumor foci, many with large clusters of ee’s (Figure 6D,F; Figure S6A). In contrast, only 20% of flies raised on lipid-depleted food contained enteroendocrine tumors, and these tumors very small (Figure 6G,H; Figure S6A). By 9 days after tumor induction, animals on control diets had large tumors, while animals fed lipid depleted food were only beginning to show tumor foci (Figure 6C, Figure S6A). To further analyze the effects of diet we induced ISC clones lacking Notch and counted clone size as a function of diet (Figure 6I-N). Nine days after induction, Notch mutant clones were smaller in animals on a lipid-depleted diet than in animals on a control or high sterol diet (Figure S6B). Interestingly, the high sterol diet affected the differentiation of tumor cells, which are normally contain ISC-like cells and ee-like cells expressing Prospero. Prospero-expressing tumor cells were suppressed on a low lipid diet, but enhanced on a high sterol diet, showing that diet can affect the nature of cell differentiation within a tumor, as well as its growth rate (Figure 6O).

## Discussion

### Dietary lipids affect Notch signaling in nutrient dependent tissues

Our experiments show that dietary lipids affect Notch signaling in both the midgut, and in other nutrient-dependent tissues such as ovarian germline and follicle cells. These effects arise from lipid-induced changes in membrane protein trafficking and stability that alter the kinetics of recycling and cellular concentration of the Notch ligand Delta and NECD (Figure 6P). The effects of dietary sterols on Notch signaling that we observed represent a widespread mechanism, because similar changes occurred in multiple sterol-requiring tissues as well as in mammalian cells. If maintained, a changed lipid environment can continue to affect Delta and NECD metabolism for an extended period. This is exactly what we observe in the ovaries of *Hr96* mutants that display high Delta levels in germ cells for a significant fraction of follicular development, showing that the Notch pathway can be chronically perturbed by changes in sterol metabolism. In the gut, if lipid-rich food is continuously provided, the lipid regions of the posterior midgut maintain relatively high sterol levels and reduced levels of Notch signaling for a significant fraction of the adult lifespan.

Not all tissues that require Notch are affected by defects in sterol metabolism. We observed no defects in wing vein patterning or bristle development in the Hr96 mutant. Even sensitive tissues have limits to the degree of modulation. When Hr96 is overexpressed, the degree of Notch signaling as measured by ee cell differentiation could not be increased further by a high sterol diet. In the absence of all Hr96 function in null mutant animals a low level of ee cells was still produced regardless of diet. This suggests that the regulation of Notch signaling by environmental lipids has natural limits that maintain at least a basal level of tissue function even under extreme circumstances. This ensures at a minimum that the tissue will still be able to respond if the nutrient environment becomes more favorable.

### Mechanism of dietary influence on Notch signaling

Delta (and NECD) localization changed dramatically on a lipid-depleted diet or in the Hr96 mutant, appearing in more vesicles and larger vesicles spread throughout the cytoplasm, and at higher levels associated with the plasma membrane. These changes, amounting to 2-3 fold increased integrated Dl immunofluorescence per cell (Figure 5), may result in small part from increased biosynthesis, since RNAseq revealed small increases in Delta mRNA (below statistical significance), but in large part from reduced Delta turnover. Increased Delta stability probably explains the major increase in Delta protein perdurance downstream from the stem cell in the enterocyte lineage. Normally only ISCs have high Delta levels, but in flies starved for cholesterol due to reduced dietary availability or Hr96 mutation, Delta remained at high levels in enteroblasts, and in differentiating enterocytes up to and including cells with 8c genomes. This equates to increased Delta persistence under lipid-starved conditions from perhaps an hour in normal enterocytes to at least the 48 hours needed for an enteroblast to develop into an 8c enterocyte. Under conditions of excess lipids or Hr96 duplication, Delta levels were reduced 2-4 fold based on fluorescence measurements.

Since the effects of dietary lipids on Notch signaling and ee production depended on the *Drosophila* LXR homolog Hr96, changes in cholesterol homeostasis in the membrane compartments of the affected cells are likely involved. Many studies have shown that a high-fat diet induces ER stress and stimulates ER associated protein degradation (ERAD). Moreover, the changes in Delta cytoplasmic distribution are exactly those expected for an alteration in stability within endosomes. General ER stress can induce ISC division (Wang et al. 2014) but we only observed increased ISC activity following lipid depletion to balance an increase in apoptosis. Rather, the substantial changes in Delta and NECD stability we observed did not affect all membrane proteins and mostly likely result from regulatory mechanisms that target these proteins specifically. Classical studies have shown that cholesterol levels can control the turnover of specific cholesterol biosynthetic enzymes like HMG-CoA reductase (Brown and Goldstein, 1980; Goldstein and Brown 1990; DeBose-Boyd, 2008; Nakanishi et al. 1988). Low cholesterol conditions stimulate ER protein trafficking of SREBP proteins (Hua et al., 1993; Wang et al., 1994; Yokoyama et al., 1993) and can result in different proteins, such as the ER regulatory protein Insig-1, being targeted for ERAD (Gong et al. 2017). Our data suggest that high sterol levels trigger Delta turnover perhaps by stimulating interaction between an Insig-like resident ER protein, the Delta protein, and an E3 ubiquitin ligase. Characterizing the precise mechanism by which lipids control the stability of Delta and NECD represents an important task for future studies.

### Dietary sensitivity of Notch signaling causes the midgut to adapt to the initial diet

Our experiments provide novel insight into how specific nutrients early in life influences intestinal structure and metabolic programming. We found that dietary cholesterol, early in adult life, plays a central role in controlling the proper proportion of ECs and ee’s in the mature *Drosophila* intestine through its effects on Notch signaling in ISCs. This provides a unique mechanism that allows the early intestine to sense available nutrients and shift the composition of the epithelia to help adapt the organism to the challenges of its nutrient environment. In mammals, after birth crypts begin to develop, new cell types emerge, and certain regional characteristics are formed (Carulli et al., 2014; Hudry et al., 2016). Many studies (Bates et al., 2006; Rakoff-Nahoum et al., 2015; Semova et al., 2012) have suggested that these postnatal effects on development may stem from seeding the intestinal flora. It is thought that the interaction between the host and the microbe underlie some of these late steps in development (Bates et al., 2006; Rakoff-Nahoum et al., 2015; Stephens et al., 2016). However, given that these later events in epithelial development occur when the animal begins to feed it is also likely that dietary nutrients play a major role in the maturation of the adult intestine.

The development and parameters of such system may not be determined by the linkage between dietary cholesterol and Notch signaling. Rather the developmental programming that causes many posterior but only a few middle and anterior ISC divisions to occur during the first 5 days after eclosion is what makes it possible for dietary lipids to modulate posterior ee number. Likewise, it may be the developmentally programmed lifetime of these ee cells that determines how long the effect will persist. As cells in the mature midgut slowly turn over, new ISC divisions will take place and the currently prevailing lipid environment that influence whether an EC or ee results. Initially lipid-deprived animals shifted to a normal diet will tend to acquire a higher proportion of ee’s and while animals raised on excess sterols shifted to a normal diet will lose ee’s with time. Dietary-induced changes in ee number that occurred quickly in the posterior of young animals, will occur gradually in the middle and anterior midgut, as we observed. Thus, our data is consistent with the idea that Notch signaling is lipid sensitive throughout much of the midgut and much of adult life, but that independently programmed ISC divisions control exactly how this connection is manifest.

How might a change in the proportion of ee’s influence the metabolic program and decisions about levels of cholesterol storage? We observed a 3-8 fold increase in 6 of the 7 recognized enteroendocrine hormone gene RNAs, consistent with the general 3-fold increase in the number of ee’s in *Hr96*^*2X*^ animals. One of the increased hormones, AstA, is closely related to allatostatin type A proteins in mammals that are ligands for the galanin receptors involved in feeding behavior and metabolic regulation (Parker and Bloom, 2012). Activation of these receptors is thought to affect food preferences, and in the brain AstA controls production of adipokinin and insulin, whose levels influence the amount of lipid stored in the fat body (Hentze et al. 2015). A smaller increase was observed in Tk, an enteroendocrine hormone that was recently implicated by ablating Tk-expressing ee’s in the negative regulation of lipid biosynthesis in the intestine and transport throughout the body (Song et al. 2014). The increases we observed in the number of Tk ee’s when lipids are in excess and reductions under lipid deprivation might correspond to homeostatic adjustments of these processes in enterocytes.

Lipid-deprivation mediated reductions in ee number persisted for at least two weeks after the animals were shifted to a rich diet. Lower ee number may explain in part our observation that 18% more whole body cholesterol was stored under these conditions, than in flies that had been raised continuously on rich medium. Persistent changes in cellular composition represents an attractive model for understanding similar metabolic adaptations in mammals. Mammalian development is strongly dependent on the mother’s nutrition, and children born to malnourished mothers often begin life with a low birth weight (Lechtig et al., 1975). These individuals are more likely to become obese later in life if nutrients are not restricted (Barbero et al., 2013; Ramakrishnan, 2004). Low nutrient exposure early in life appears to trigger metabolic changes that enhance nutrient scavenging by the developing fetus. After birth, these resets become a permanent state that puts individuals at risk for metabolic syndromes in a normal nutritional environment (Barbero et al., 2013; Reusens et al., 2011).

### Other regions of the digestive system may adapt to the dietary levels of different nutrients

Nutrients in addition to cholesterol appear to be absorbed and metabolized in unique regions of the intestine (Buchon et al., 2013; Marianes and Spradling, 2013). Like cholesterol, these nutrients may establish a feedback mechanism where a region of the intestine that absorbs a specific nutrient class senses that nutrient and changes the cellular composition that helps the organ adapt their metabolism and utilization of that nutrient. For example, previous work has shown that HNF4 functions as a fatty acid sensor that controls gluconeogenesis and fatty acid oxidation (Palanker et al., 2009). Interestingly, other studies have shown that HNF4s also plays important roles in the differentiation and development of the tisssues in the digestive tract (Dhe-Paganon et al., 2002; Hayhurst et al., 2001). Other nuclear receptors such as ERR have been shown mediate growth and tissue differentiation in the pancreas and cardiac muscle by controlling changes in glycolysis and mitochondrial oxidative metabolism that are essential for the development and function of mature tissues(Alaynick et al., 2007; Tennessen et al. 2011; Yoshihara et al., 2016). Thus, the types of adaptation described here for dietary cholesterol, may be widespread during the final differentiation of multiple regions of the digestive system.

### Dietary sterols, Notch signaling and cancer

The connection between sterol metabolism and Notch signaling is not only important for metabolic adaptation but has potentially strong implications for cancer etiology and therapeutics. Mutations in the Notch signaling pathway have been associated with cancer deriving from many tissues in the digestive tract including the liver, pancreas and the colon. Furthermore, Notch signaling has been a common therapeutic target for these types of cancer. However, drugs and therapies targeting Notching signaling commonly have severe side effects throughout the body complicating their use with patients. We found that stimulating reverse cholesterol transport using LXR agonists in human embryonic kidney cells (HEK293) caused a dramatic and highly reproducible decrease in Delta-like-ligand 4 protein levels. This suggests that sterol metabolism may have a highly conserved role in the regulation of Notch signaling that could be exploited for therapeutic purposes. For example, sterol regulating drugs such as LXR agonists may provide a way to alter Notch signaling in a tissue-directed manner for adjuvant therapy and as a means to reduce cancer risk in genetically susceptible patients. Inhibiting Delta like ligand 4 (DLL4) is sufficient to reduce tumor growth and initiation in mice and humans (Hoey et al., 2009).

Numerous studies have shown that consuming a high-fat diet increases susceptibility to several specific types of cancer (Breitkreutz et al., 2005; Huang et al., 2012; Mattisson et al., 2004; Tang et al., 2012). High-fat diets have been proposed to increase cronic inflammation in tissues including liver and intestine (Ding et al., 2010; Khalil et al., 2010; Miyagi et al., 2010; Shrestha et al., 2013; van der Heijden et al., 2015; Wu et al., 2013). Moreover, a high fat diet promotes stemness and tumorigenicity in intestinal organoid cultures (Beyaz et al. 2016), however, the lipids causing this effect and the underlying mechanisms remain unclear. Our study provides a specific mechanism to explain these connections. Large quantities of dietary lipids, particularly sterols, may directly inhibit tumor suppressor pathways, such as Notch signaling, and increase cancer susceptibility in a manner that synergizes with inflammation. In contrast, consuming a low-fat diet has been suggested to protect against multiple types of cancer. Consistent with this idea we found that feeding flies a diet low in dietary sterols suppresses enteroendocrine tumor growth and tumor size. Moreover, we found that the relative proportion of tumor cells expressing a differentiation marker, Prospero, was strongly affected by diet. Taken together our work suggests that dietary lipid levels play a central role in dictating cancer risk for digestive track tumors and that dietary intervention in conjunction with the use of cholesterol modulating drugs may provide an effective strategy for reducing cancer risk in genetically predisposed patients.

### Evolutionary origin of nutrient regulation of development

The ability of cells to maximize their utilization of critical resources such as environmental sterols is critically important to both unicellular organisms as well as metazoans. Mechanisms for optimizing cholesterol uptake, biosynthesis, transport and excretion evolved early and are widely shared throughout unicellular and metazoan animals and plants. Perhaps because lipid uptake inherently involves endocytosis and vesicle trafficking, regulatory mechanisms controlling lipid utilization in animals are centered on the stability of key regulatory endosomal membrane proteins (Brown and Goldstein, 1990). Since lipids are constituents of endosomal membranes their uptake is likely to affect the physical properties of endoplasmic reticulum membranes which lead to binding to sterol sensing domains in proteins such as Insig that can bring ubiquitin ligases in contact with effector membrane targets.

The experiments reported here suggest that this ancient system extends beyond the control of nutrient homeostasis and metabolism to include development. The intestine is perhaps the most directly important tissue for bringing environmental nutrients into balance with organismal requirements, both in the short term and by anticipating future needs. The ovary is the most important tissue for ensuring the success of the next generation. The Notch signaling pathway is central to the differentiation of cells downstream from stem cells whose presence in appropriate numbers can contribute to the success of these tasks. We have found that endosomal trafficking of the key Notch membrane proteins Notch and Delta is lipid sensitive, like other key membrane regulators of nutrient utilization and metabolism. Alterations in the levels of these proteins affects Notch signaling and cell fate specification downstream from intestinal, germline and follicle cell stem cells. These findings represent a particularly clear example where the nutrient environment exerts specific, direct, and measurable effects on specific proteins by well understood mechanisms that mediate cell renewal, and cell type homeostasis. Further characterizing these connections will help reveal how environmental nutrients can positively impact tissues that have become stressed due to age, disease or the presence of cancer.

## Acknowledgements

We thank Steve Deluca, Ethan Greenblatt, Chenhui Wang and members of the Spradling lab for discussion and valuable comments on the manuscript. RO is a student in the Cellular, Molecular and Developmental Biology graduate program of Johns Hopkins University. MS was a fellow of the Jane Coffin Childs Memorial Fund.

## Author contributions

RO, MS and ACS designed experiments and wrote the manuscript. RO and MS performed research.

